# Dengue modeling in rural Cambodia: statistical performance versus epidemiological relevance

**DOI:** 10.1101/208876

**Authors:** Clara Champagne, Richard Paul, Sowath Ly, Veasna Duong, Rithea Leang, Bernard Cazelles

## Abstract

Dengue dynamics are shaped by the complex interplay between several factors, including vector seasonality, interaction between four virus serotypes, and inapparent infections. However, paucity or quality of data do not allow for all of these to be taken into account in mathematical models. In order to explore separately the importance of these factors in models, we combined surveillance data with a local-scale cluster study in the rural province of Kampong Cham (Cambodia), in which serotypes and asymptomatic infections were documented. We formulate several mechanistic models, each one relying on a different set of hypotheses, such as explicit vector dynamics, transmission via asymptomatic infections and coexistence of several virus serotypes. Models are confronted with the observed time series using Bayesian inference, through Markov chain Monte Carlo. Model selection is then performed using statistical information criteria, but also by studying the coherence of epidemiological characteristics (reproduction numbers, incidence proportion, dynamics of the susceptible class) in each model. Considering the available data, our analyses on transmission dynamics in a rural endemic setting highlight both the importance of using two-strain models with interacting effects and the lack of added value of incorporating vector and explicit asymptomatic components.

## 1 Introduction

Dengue is a vector-borne viral disease transmitted by *Aedes* spp. caused by any of four dengue virus (DENV) serotypes. Infection can result in a flu-like illness, and sometimes potentially lethal complications called Dengue Hemorrhagic Fever (DHF) and Dengue Shock Syndrome (DSS), although a significant proportion are subclinical or asymptomatic, causing insufficient discomfort for clinical presentation [1]. Dengue is ubiquitous in the tropics and the subtropics, particularly in Southeast Asia, the Pacific and the Americas [2]. The World Health Organization (WHO) considers that dengue is a major public health issue worldwide, with four billion people in 128 countries exposed to the dengue virus [3, 4], an estimated 390 million infections every year and about 50-100 million symptomatic cases worldwide and a high disease burden [5, 6]. Nowadays, there are more cases of dengue worldwide than any other arboviral disease [7, 8, 9].

The value of mathematical models and associated statistical tools for investigating public health policy questions has long been recognized and has provided insights into their transmission and control for more than one hundred years [10, 11]. It is important, however, to adapt them as much as possible to a specific setting, in order to derive appropriate public health recommendations and accurately generate the key parameters using estimation tools, so that they can produce realistic conclusions, in accordance with the observed data.

Dengue dynamics are shaped by the complex interplay between many factors associated with the mosquito vector and human hosts and their interactions with the virus. Hitherto, the exploration of dengue dynamics has focused on the urban setting, where the incidence of dengue is highest [12, 13, 14, 15]. Few studies have been carried out in rural settings [16, 17, 18], despite growing evidence that rural dengue is an increasing problem. Guha-Sapir and Schimmer [19] observed shifts in modal age, rural spread, and social determinants of dengue susceptibility, with major implications for health services. Muhammad Azami et al. [20] observed similar dengue seroprevalence rates between urban and rural samples, showing that dengue is not confined to urban areas in Malaysia. Chareonsook et al. [21] showed that DHF in Thailand, which was originally thought to be an urban disease, has spread to most areas of Thailand, and is now more common in rural than urban areas and studies suggest that rural dengue incidence can surpass urban and semi-urban communities within the same region [22, 23]. In addition, several studies have stressed that rural settings play an important role in the timing of dengue epidemics in Southeast Asia, with the seasonal dengue waves typically arriving later in major urban centers [24, 25, 26].

In this study, we combine two datasets from rural Cambodia that provide information on different key factors. We contrast and compare several mechanistic models, incorporating differing levels of complexity with respect to vector dynamics, coexistence of several virus strains, and transmission via asymptomatic infections. Models are adapted to the observed time series using Bayesian inference, through Markov Chain Monte Carlo (MCMC) and compared in light of the data, using statistical indicators to identify the best model [27, 28, 13, 29]. In addition, we also analyze the epidemiological coherence of the estimated models in simulations. Critically, we do not merely focus on the observed infected individuals but also on other compartments, such as the susceptible class of individuals. By comparing these models, we try to find a realistic but parsimonious way of modeling dengue epidemics in rural Cambodia. The best model may then be used in the study of intervention scenarios or in comparative analyses with other settings. For instance, it could be readily expanded to understand the potential impact of different vaccination strategies in rural settings.

## 2 Methods

### 2.1 Data

#### 2.1.1 Study area

Kampong Cham province is a densely populated rural province 120km northeast from the capital Phnom Penh. Dengue is endemic and strongly seasonal (cf. Figure 1), with outbreaks occuring every year from June to September, during the rainy season. The four virus serotypes co-circulate, even though one usually dominates the three others for about 3 to 5 years. We used two different datasets reporting dengue cases in the province: the results of a punctual study conducted in a 30km radius around the city of Kampong Cham (DENFREE data), and the national surveillance data (NDSS data) in the four districts comprising the DENFREE study area (Kampong Cham, Kampong Siem, Prey Chhor and Tboung Khmum, with the administrative divisions of 2012-2013).

**Figure 1.**
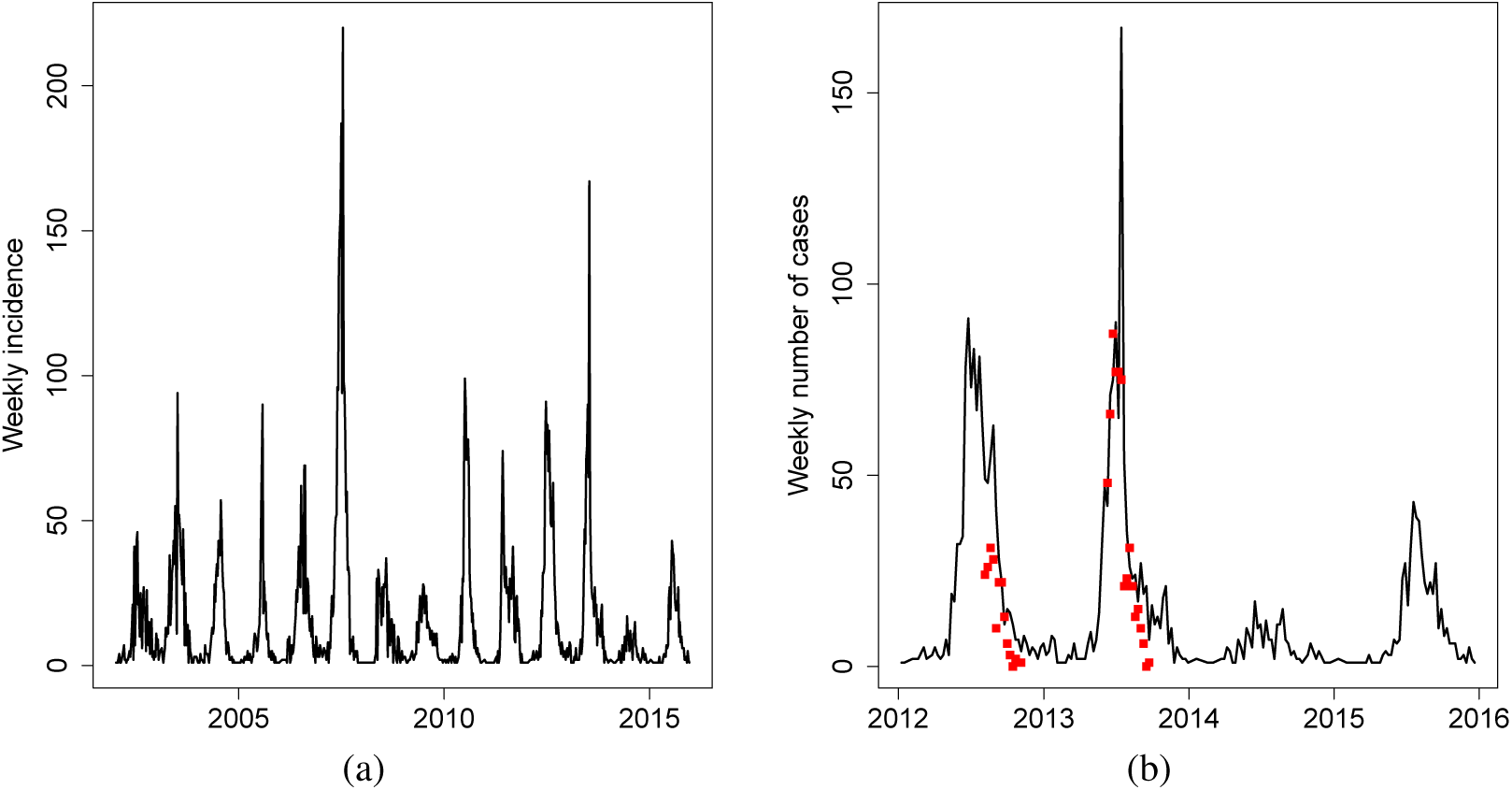
**a) NDSS data.** Weekly number of cases in the four districts of Kampong Cham province. **b) NDSS and DENFREE data.** The black line gives the series of NDSS cases per week in the four districts of the Kampong Cham province. The red dots are the DENFREE cases per week in the DENFREE study area.

#### 2.1.2 DENFREE Data

The DENFREE study took place in the Kampong Cham region during the 2012 and 2013 outbreaks. Patients with acute dengue-like illness were enrolled in three hospitals in the Kampong Cham province and DENV infection was confirmed by qRT-PCR. Positive DENV cases were considered as index cases, and an outbreak investigation was initiated in their neighbourhood, in order to detect asymptomatic or mildly symptomatic cases. The study protocol is extensively detailed in Duong et al. [30].

We used the series of the total number of cases per week (index cases and outbreak investigation cases) between August 6th (first week when more than 100 people were tested for dengue in outbreak investigation) and November 5th 2012 and between June 10th (first week with outbreak investigation) and September 23rd 2013. We also restricted the study to children under 15 years old for two major reasons : most of the reported dengue cases were in this age class, and it allowed a comparison with other dengue reporting systems in Cambodia, which are mainly done at paedriatric hospitals. Information on the serotype responsible for infection, and symptomatic/asymptomatic status of the patients were available.

**Table 1.** **Number of cases (index cases and community cases) under 15 years old, in the DENFREE study.** Cases are restricted to the period 6th Aug - 5th Nov 2012 and 10th Jun - 23rd Sept 2013. During the whole study, 236 symptomatic cases were collected in 2012 and 574 in 2013. Serotype was unknown for 5 cases in 2012. One case in 2013 was coinfected with DENV-1 and DENV-2 and was not included in models with two strains (serotypes).

| Children under 15 years old | 2012 | 2013 |
| --- | --- | --- |
| **Observed symptomatic cases** | 189 | 571 |
| Denv-1 | 183 | 448 |
| Denv-2, Denv-3, Denv-4 | 1 | 122 |
| **Observed asymptomatic cases** | 4 | 28 |
| **Children tested in the community** | 1722 | 4119 |

#### 2.1.3 NDSS Data

Because the DENFREE data covers only a relatively short period of time, surveillance data were added to improve the estimations. Surveillance of dengue is conducted at the national level in Cambodia, through the National Dengue Surveillance system (NDSS)[31, 24], involving the paediatric departments of several hospitals throughout the country. Diagnosis is done clinically and only a small fraction of the cases are confirmed serologically. Because of the co-circulation of other flaviviruses (Chikungunya, Japanese Encephalitis) and the relative non-specificity of symptoms, clinical mis-diagnosis may be frequent. Since surveillance is carried out in paediatric departments, only cases among children under 16 years old are reported.

We selected all the cases under 15 years old in the four districts involved in the DENFREE study between January 2002 and December 2015 and aggregated them per week (cf. Figure 1). In this area, on average, 770 cases under 15 years old are reported per year (maximum 1985 cases in 2007, minimum 209 cases in 2014). We used data from 2002 to 2013 for estimations, and data for 2014 and 2015 as the test set. As displayed in figure 1, NDSS and DENFREE data have similar dynamics.

**Table 2.** **Comparison of both datasets**

| DENFREE | NDSS |
| --- | --- |
| Observations in 2012-2013 | Observations in 2002-2015 |
| No observation during inter-epidemic period, dataset starts during the epidemic peak | Observations all year round |
| Laboratory confirmation | Clinical diagnosis |
| Observation of non hospitalized cases including some asymptomatic infections | Hospital cases only |
| Clustered collecting process | Stable reporting process over time |
| An observation rate can be estimated | Unknown observation rate |
| Known serotype | Unknown serotype |

#### 2.1.4 Population

We take as the reference population (N=161391) the number of children below 15 years old in four districts of the Kampong Cham province (Kampong Cham, Kampong Siem, Prey Chhor and Tboung Khmum, with the administrative divisions of 2012-2013) according to 2008 National Census [32]. Since the DENFREE study was conducted in a subpart of this area, we calculated the total population for the DENFREE study (n=65208) as the sum of the population of children under 15 years old in all the villages investigated in either 2012 or 2013 [32].

### 2.2 Models

All model parameters are defined in the figures captions and in Table 3.

**Table 3.**
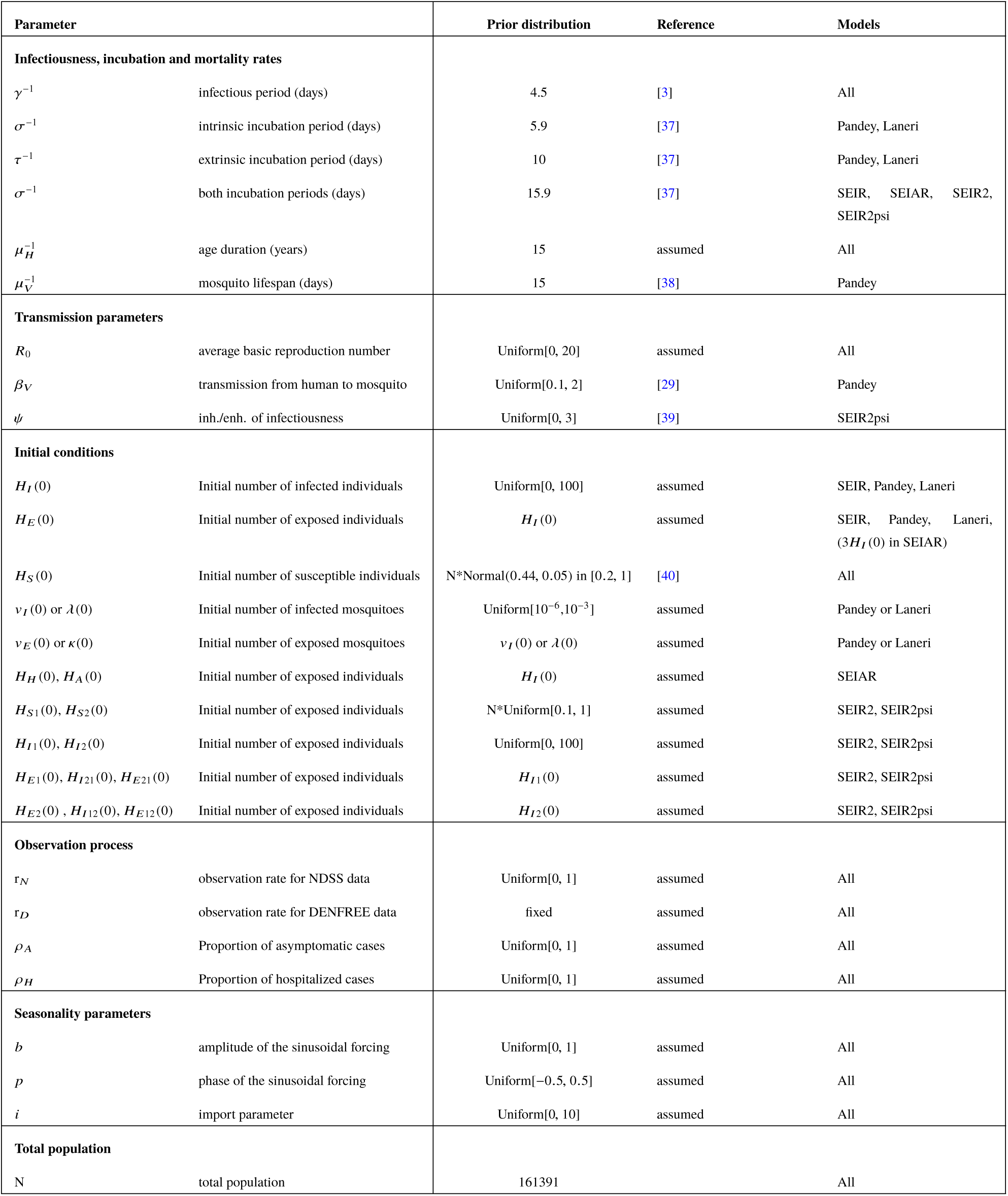
**Prior distributions of parameters**. "Uniform[0,20]" indicates a uniform distribution in the range [0,20]. "Normal(4.5,0.1) in [4,5]" indicates a normal distribution with mean 4.5 and standard deviation 0.1, restricted to the range [4,5].

#### 2.2.1 One-strain models

We take a Susceptible-Exposed-Infected-Recovered (SEIR) model as the simplest model (cf. Figure 2). In this model, the basic reproduction number, i.e. the number of secondary human infections resulting from the introduction of a single infected individual in an entirely susceptible population, is 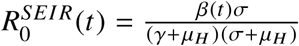.

**Figure 2.**
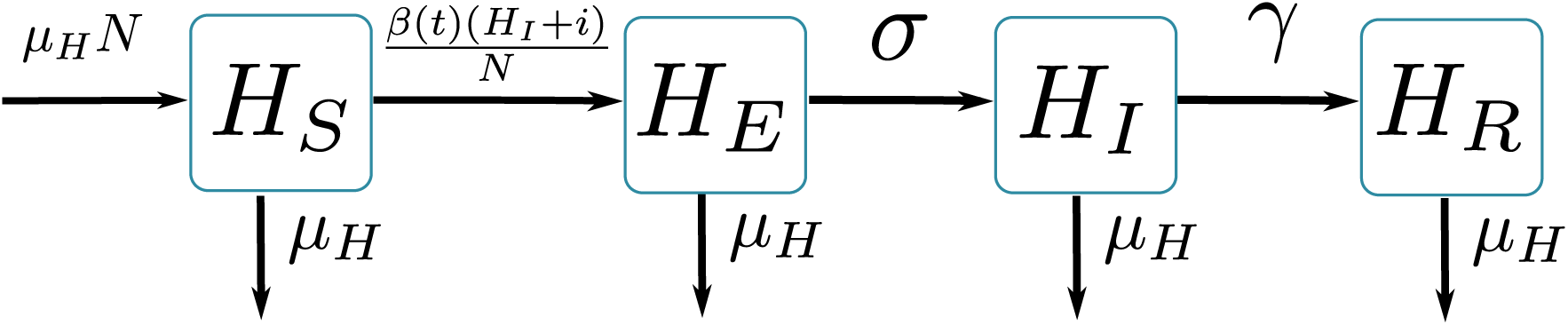
Graphical representation of SEIR model. *H*_*S*_ susceptible individuals; *H*_*E*_ infected (not yet infectious) individuals; *H*_*I*_ infectious individuals; *H*_*R*_ recovered individuals; **β**(*t*) is the transmission parameter; **σ** is the rate at which *H*_*E*_-individuals move to the infectious class *H*_*I*_; infectious individuals (*H*_*I*_) then recover at rate **γ**; individuals leave the children population at rate **µ**_*H*_. *H*_*S*_ + *H*_*E*_ + *H*_*I*_ + *H*_*R*_ = *N*.

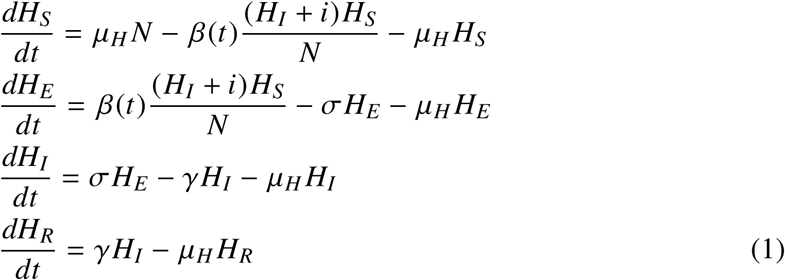

This model is compared with two other models that include the mosquito vector transmission components. In the first one, derived from Pandey et al. [29], the vector is modelled explicitly with three compartments (Susceptible-Exposed-Infected) (cf. Figure 3). In the second one, derived from Laneri et al.[33], the vector is modelled implicitly as an external force of infection including two stages, latent (*κ*) and current (*λ*) (cf. Figure 4). We derived *R*_0_ for each model as 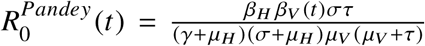 and the estimation 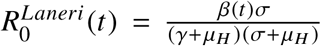 [34]. In order to compare these models with the non-vector models, we considered the same definition (i.e. the number of secondary human infections resulting from the introduction of a single infected individual in a entirely susceptible population), and not the reproduction ratio per generation provided through the use of the next generation matrix.

**Figure 3.**
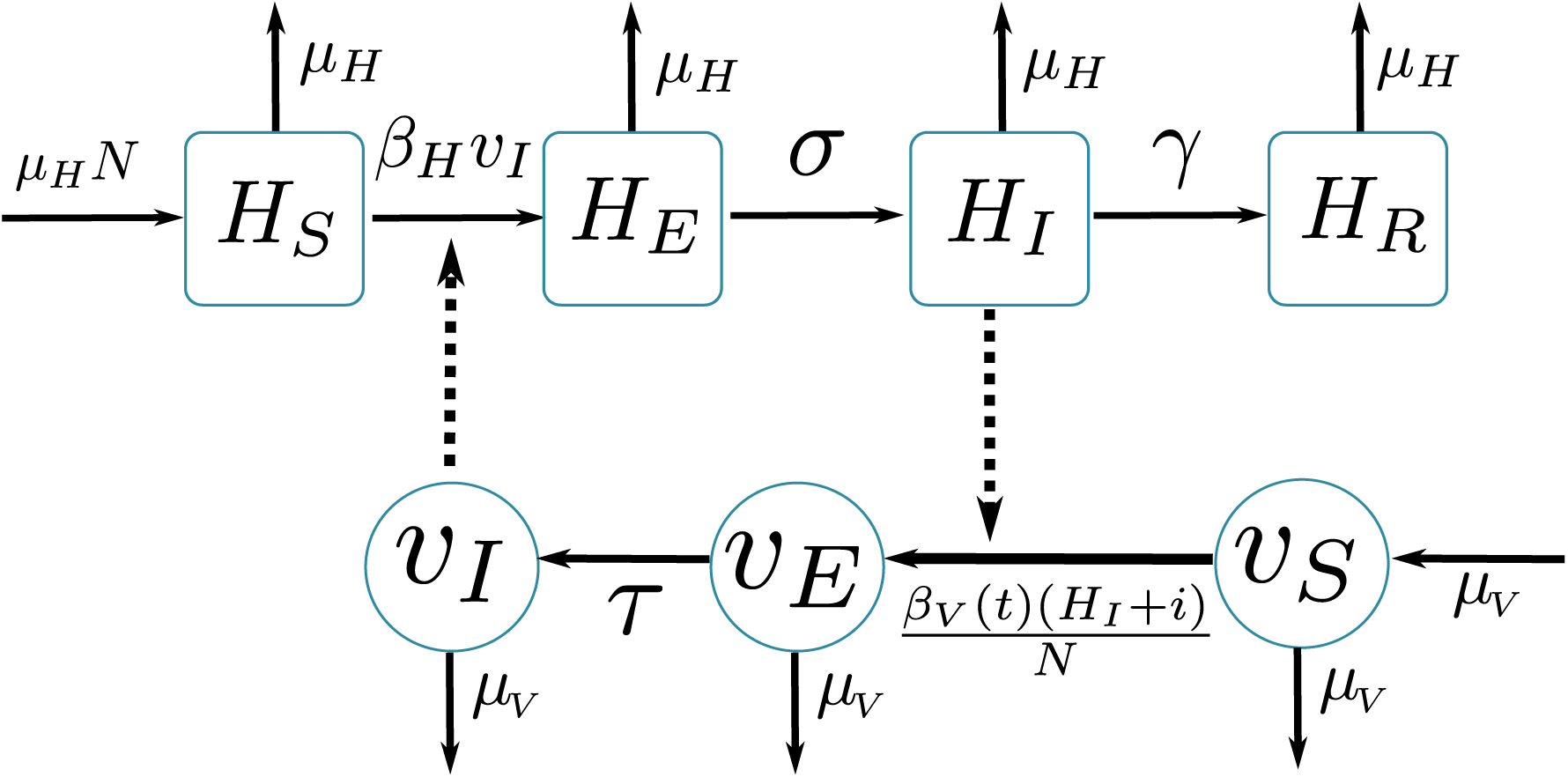
Graphical representation of Pandey model [29]. Squared boxes and circles correspond respectively to human and vector compartments. Plain arrows represent transitions from one state to the next. Dashed arrows indicate interactions between humans and vectors. *H*_*S*_ susceptible individuals; *H*_*E*_ infected (not yet infectious) individuals; *H*_*I*_ infectious individuals; *H*_*R*_ recovered individuals; **β**_*H*_ is the transmission parameter from vector to human; **σ** is the rate at which *H*_*E*_ -individuals move to the infectious class *H*_*I*_; infectious individuals (*H*_*I*_) then recover at rate *γ*; individuals leave the children population at rate **µ**_*H*_; *H*_*S*_ + *H*_*E*_ + *H*_*I*_ + *H*_*R*_ = *N*; *v*_*S*_ proportion of susceptible vectors; *v*_*E*_ proportion of infected (not yet infectious) vectors; *v*_*I*_ proportion of infectious vectors; *β*_*V*_(*t*) is the transmission parameter from human to vector; *τ* is the rate at which *v*_*E*_ -vectors move to the infectious class *v*_*I*_; vectors die at rate *µ*_*V*_.

**Figure 4.**
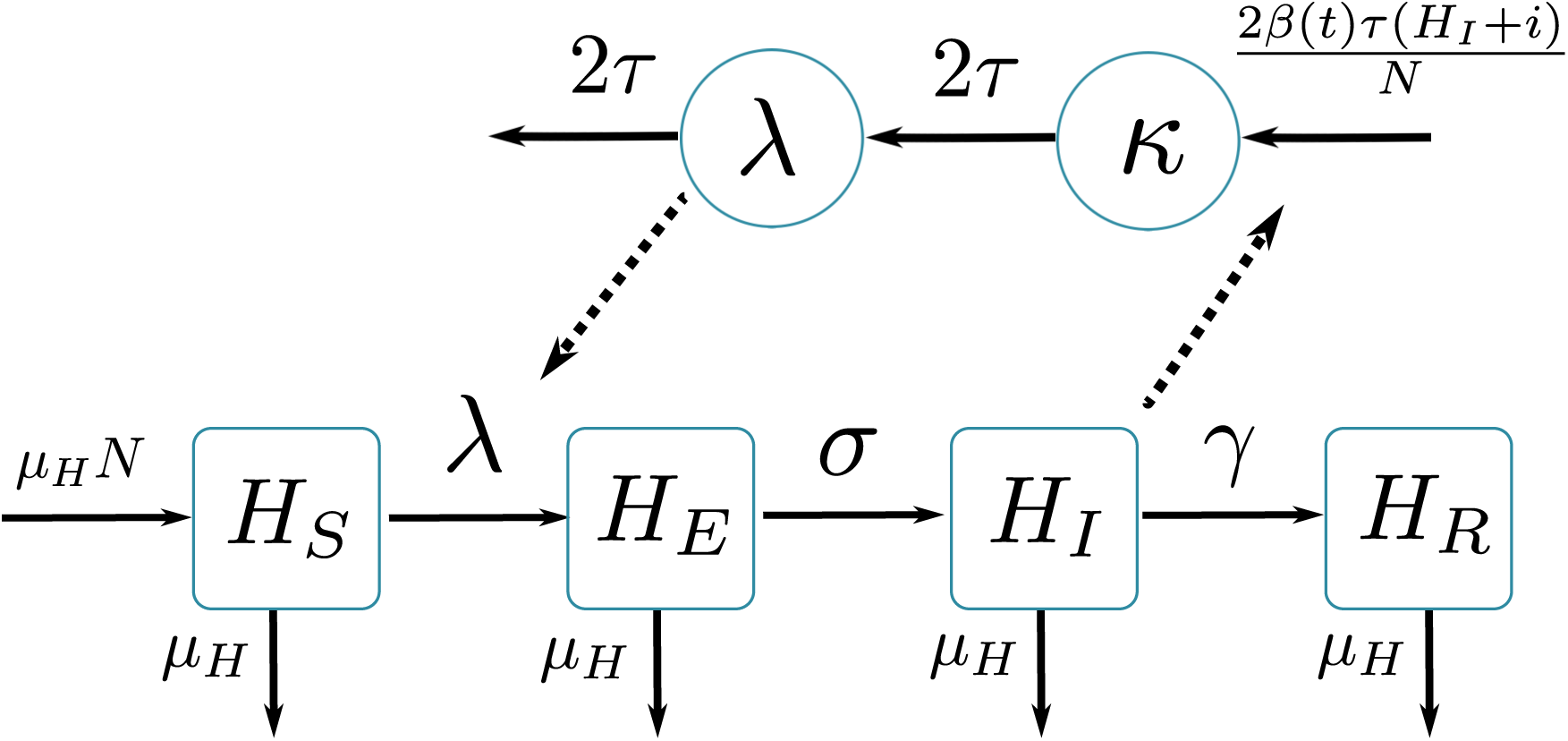
Graphical representation of Laneri model [33]. Squared boxes and circles correspond respectively to human and vector compartments. Plain arrows represent transitions from one state to the next. Dashed arrows indicate interactions between humans and vectors. *H*_*S*_ susceptible individuals; *H*_*E*_ infected (not yet infectious) individuals; *H*_*I*_ infectious individuals; *H*_*R*_ recovered individuals; *σ* is the rate at which *H*_*E*_ -individuals move to the infectious class *H*_*I*_; infectious individuals (*H*_*I*_) then recover at rate *γ*; individuals leave the children population at rate *µ*_*H*_; *H*_*S*_ + *H*_*E*_ + *H*_*I*_ + *H*_*R*_ = *N*; implicit vector-borne transmission is modelled with the compartments *κ* and *λ*; *λ* current force of infection; *κ* latent force of infection reflecting the exposed state for mosquitoes during the extrinsic incubation period; *β*(*t*) is the transmission parameter; *τ* is the transition rate associated with the extrinsic incubation period.

The equations describing the Pandey model are:

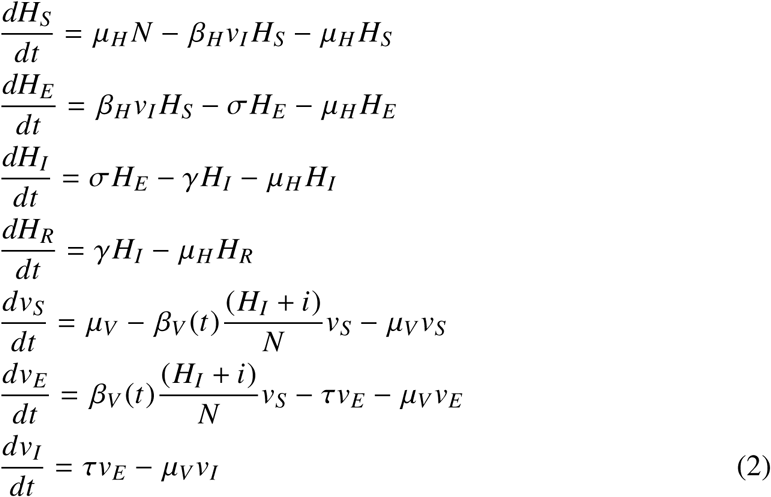

where *v*_*s*_ is the proportion of susceptible mosquitoes, *v*_*E*_ the proportion of exposed mosquitoes, and *v*_*I*_ the proportion of infected mosquitoes.

The equations describing the Laneri model are:

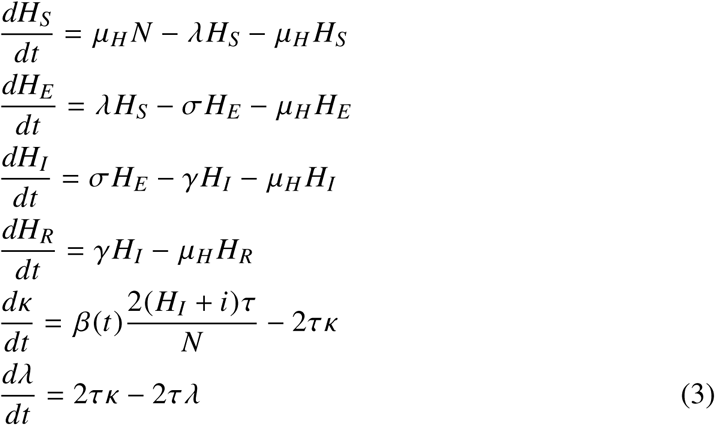

#### 2.2.2 Model with explicit asymptomatic individuals (SEIAR)

We also consider a model in which asymptomatic infections are explicitly taken into account in the transmission process (cf. Figure 5). In this model, we assume that, after the incubation period, there are three possible manifestations of the disease: asymptomatic (H_*A*_), mildly symptomatic not requiring hospitalization (H_*I*_) and hospitalized cases (H_*H*_). Asymptomatic cases are defined in the dengue study as asymptomatic or pauci-symptomatic (presence of other symptoms not being sufficient to classify as symptomatic). Hospital cases are defined as NDSS cases (reported by the surveillance system in hospitals). We assume that symptomatic DENFREE cases are either (H_*I*_) or (H_*H*_). We also assume that asymptomatic cases transmit the disease as much as symptomatic cases, as recently shown [30], and therefore, 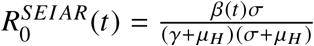.

**Figure 5.**
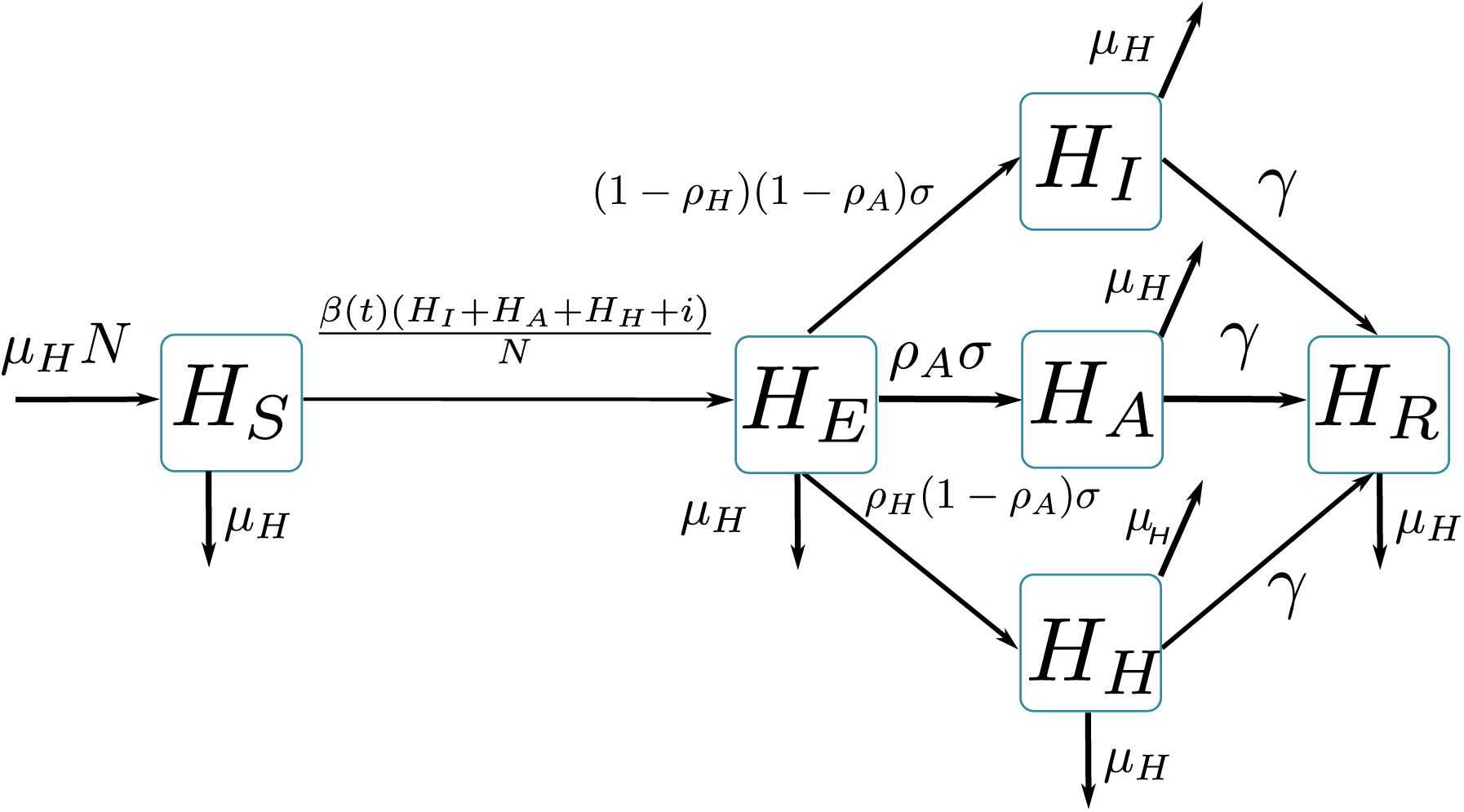
Graphical representation of SEIAR model. *H*_*S*_ susceptible individuals; *H*_*E*_ infected (not yet infectious) individuals; *H*_*A*_ asymptomatic infectious individuals; *H*_*I*_ mildly symptomatic infectious individuals; *H*_*H*_ hospitalized infectious individuals; *H*_*R*_ recovered individuals; *β*(*t*) is the transmission parameter; *σ* is the rate at which *H*_*E*_-individuals move to the infectious classes *H*_*I*_, *H*_*A*_ and *H*_*H*_; a proportion *ρ*_*A*_ of *H*_*E*_-individuals do not show symptoms during the infectious period; a proportion *ρ*_*H*_ of symptomatic individuals go to hospital; infectious individuals (*H*_*I*_,*H*_*A*_,*H*_*H*_) then recover at rate *γ*; individuals leave the children population at rate *µ*_*H*_. *H*_*S*_ +*H*_*E*_ +*H*_*A*_+*H*_*I*_ +*H*_*H*_ +*H*_*R*_ = *N*.

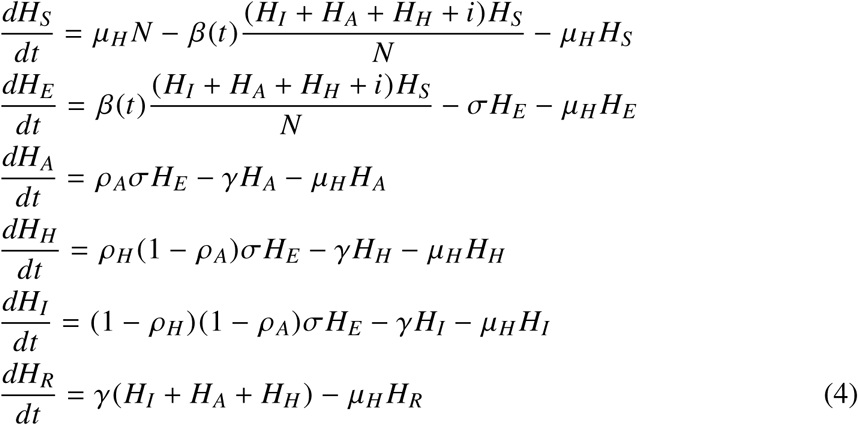

#### 2.2.3 Model with two virus serotypes

In the 2012 and 2013 epidemics, DENV-1 was highly dominant : the three other serotypes represented less than 1% of the cases reported in the DENFREE study in 2012 and about 20% in 2013 (cf. Table 1). Therefore, a two-strain model is also studied, in which we separate DENV-1 cases from DENV-2, DENV-3 and DENV-4 combined (cf. Figure 6). For simplicity and parsimony in the number of parameters, the two strains share the same parameter values. We first assume both strains to be independent (*ψ* = 1 in equation 5, called SEIR2 model). In this context, the reproduction numbers for each strain are equal, 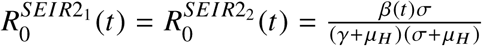.

**Figure 6.**
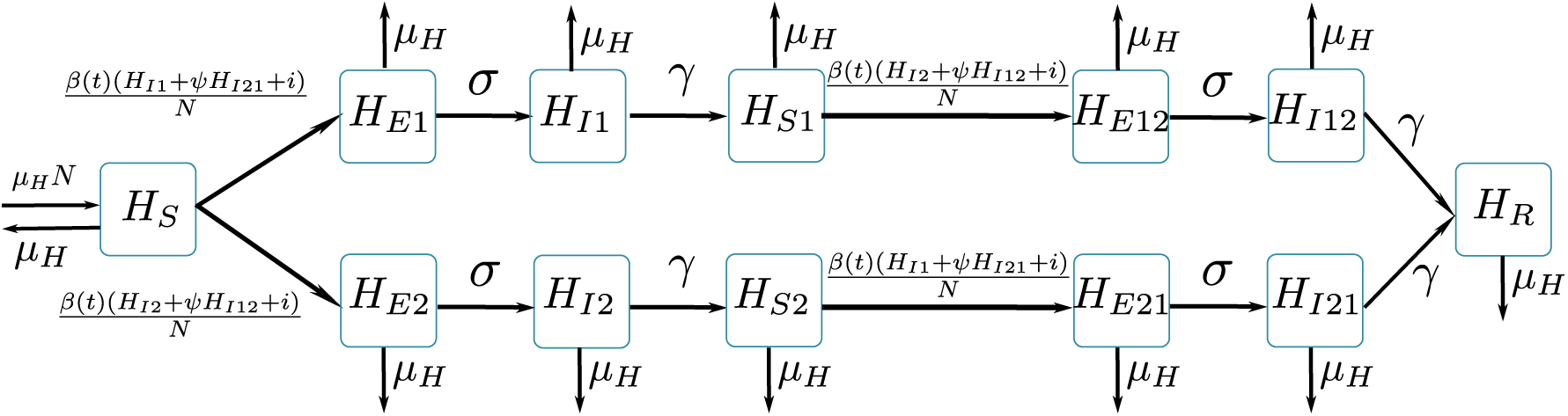
Graphical representation of SEIR2 models. *H*_*S*_ individuals susceptible to both strains; *H*_*E*1_ (resp. *H*_*E*2_) individuals infected (not yet infectious) to strain 1 (resp. strain 2); *H*_*I*1_ (resp. *H*_*I*2_) individuals infectious to strain 1 (resp. strain 2); *H*_*S*1_ (resp. *H*_*S*2_) individuals immune to strain 1 only (resp. strain 2); *H*_*E*12_ (resp. *H*_*E*21_) individuals (not yet infectious) with a secondary infection to strain 2 (resp. strain 1); *H*_*I*12_ (resp. *H*_*I*21_) infectious individuals with a secondary infection to strain 2 (resp. strain 1); *H*_*R*_ individuals immune to both strains; *β*(*t*) is the transmission parameter; *σ* is the rate at which exposed individuals move to the infectious class; infectious individuals then recover at rate *γ*; *ψ* is the change in infectivity for secondary infected individuals in SEIR2psi model (in SEIR2 model, *ψ* = 1); individuals leave the children population at rate *µ*_*H*_. *H*_*S*_ + *H*_*E*1_ + *H*_*E*2_ + *H*_*I*1_ + *H*_*I*2_ + *H*_*S*1_ + *H*_*S*2_ + *H*_*E*12_ + *H*_*E*21_ + *H*_*I*12_ + *H*_*I*21_ + *H*_*R*_ = *N*.

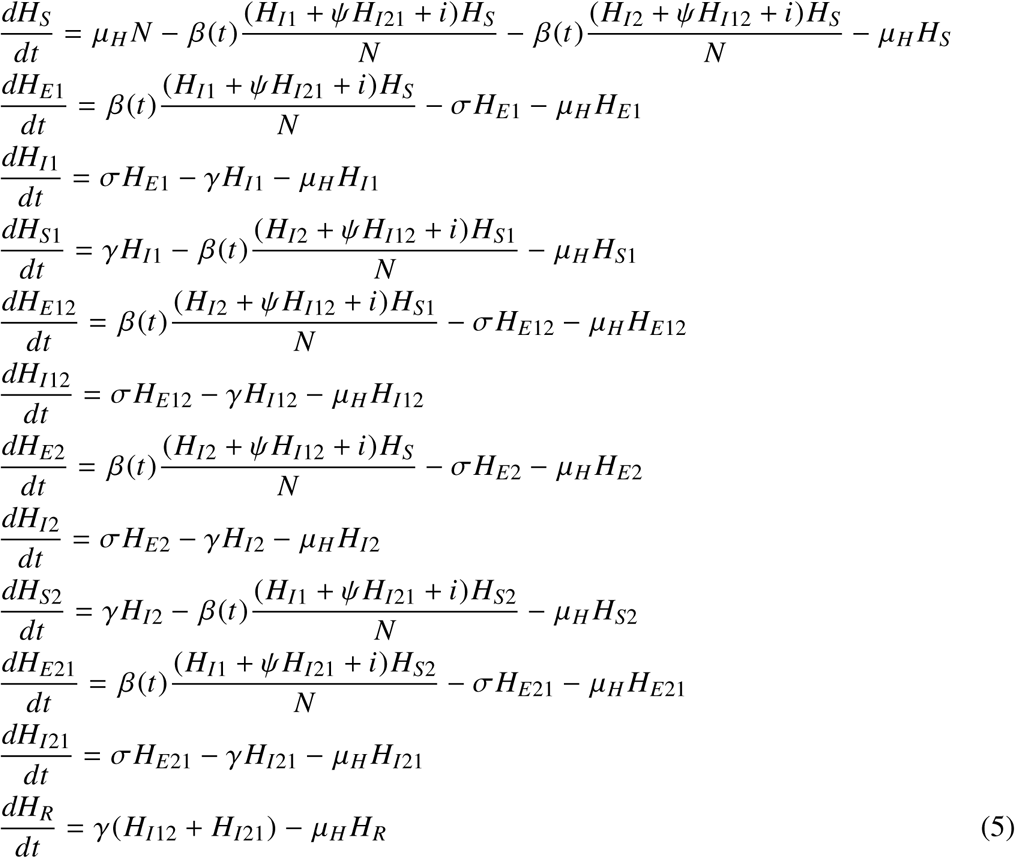

We also considered another version of the model including interaction between strains, in order to reflect the fact that secondary infection with a heterologous serotype leads more often than primary infection to severe manifestations of the disease [7]. In our model (called SEIRpsi model), primary and secondary infections differ in infectiousness, through a parameter *ψ* [35]. This parameter is estimated between zero and three: values superior to 1 correspond to transmission cross-enhancement (because of higher virus titers during secondary infections [35]) and values inferior to 1 suggest a lower infectivity for secondary infected individuals (for example because they are hospitalized and less in contact with the population [36]). As in Ferguson et al. [35], we define 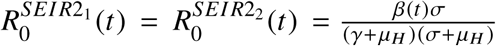 the basic reproduction number for each strain.

In the DENFREE data, DENV-1 cases are assumed to be either *H*_*I*1_ or *H*_*I*21_ and DENV-2/ DENV-3/ DENV-4 cases *H*_*I*2_ or *H*_*I*12_. In the NDSS data, observed cases are assumed to be *H*_*I*_1, *H*_*I*_2, *H*_*I*_12 or *H*_*I*_21.

#### 2.2.4 Seasonality

All models include seasonality through the use of a time-varying transmission parameter 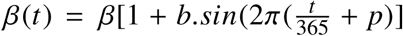, according to a sinusoidal function whose phase *p* and amplitude *b* are estimated.

We also assume that a constant number of cases *i* are imported.

#### 2.3 Prior distributions

The prior distributions are listed in Table 3.

Dirac priors based on the literature were used for the durations of infectiousness and incubation, as well as the mortality rates. In models without vectorial transmission, the incubation period is assumed to be the sum of the extrinsic (in mosquito) and intrinsic (in human) incubation periods, to reflect the generation time of the disease. For transmission parameters, we used wide weakly informative priors.

#### 2.3.1 Initial conditions

The initial number of infected individuals is assumed to be equal to the number of exposed individuals and to be lower than 100, as the model starts in January, during the epidemic trough. Except for the initial proportion of susceptibles, all priors on initial conditions are uniform distributions.

The initial proportion of susceptibles is an influential parameter on the model outputs. It is highly correlated to the transmission parameter *β* (and therefore to the basic reproductive number), which makes it difficult to estimate them both. An informative gaussian prior was therefore used on *H*_*S*_(0). To date, no large scale seroprevalence study is available for Cambodia, and we relied on a study conducted among schoolchildren in rural Vietnam [40], which we considered as the closest setting to be compared with Kampong Cham. We extrapolated their results on schoolchildren (7 to 14 years old) to a 1-15 years old population as follows, where *S*_*λ*_ is the proportion of susceptibles among 1-15 year-old children (using their estimation *λ* = 0.117):

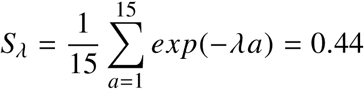

*S*_*λ*_ is used as the mean of the gaussian prior, and the standard deviation is fixed at 0.05.

#### 2.3.2 Observation rate

Using information on the sampling scheme, we calculate an observation rate on DENFREE data (cf. Table 4). We assume that index cases are all reported and that the observation rate for community cases equals the ratio of people tested over the population of the area. We then extrapolate this observation rate to the total population of the four districts.

**Table 4.** **Calculation of the observation rate for DENFREE data.** These calculations include all cases under 15 years old in the DENFREE study (18th Jun - 24th Dec 2012 and 3rd Jun - 23rd Sep 2013).

| Children < 15 years old | 2012 |  | 2013 |  |
| --- | --- | --- | --- | --- |
|  | Symptomatic | Asymptomatic | Symptomatic | Asymptomatic |
| Population in investigated villages (n) | 65,208 |  | 65,208 |  |
| Population in the 4 districts (N) | 161,391 |  | 161,391 |  |
| Index cases (a) | 151 | 0 | 376 | 0 |
| Observation rate for index cases | 1 | . | 1 | . |
| Children tested in communities (b) | 1722 |  | 4119 |  |
| DENV positive in communities (c) | 85 | 5 | 198 | 28 |
| Observation rate for community cases (d=b/n) | 0.0264 |  | 0.0632 |  |
| Extrapolated number of community cases (e=c/d) | 3219 | 189 | 3135 | 443 |
| Extrapolated total number of cases (f=a+e) | 3370 | 189 | 3511 | 443 |
| Observed total number of cases (g=a+c) | 236 | 5 | 574 | 28 |
| **Observation rate ($r_s=(g/f)*(n/N)$ and $r_a=b/N$)** | **0.0283** | **0.0107** | **0.0661** | **0.0255** |

Different observation rates are used in the SEIAR model. For DENFREE cases, the observation rate is assumed known and relies on the fact that the entire population was not tested during the study. For NDSS data, the observation rate is interpreted as the proportion of hospitalized cases (which assumes that all hospitalized cases go through surveillance and neglects the presence of private hospitals or non reports from hospitals).

### 2.4 Estimation

Models are performed in deterministic framework and estimations are made using random walk Metropolis Hastings. SSM software [41] is used for simulations and calculations. The observation model is negative binomial with the dispersion parameter fixed at 0.1 [42]. In order to initialize the starting point of MCMC chains, we ran a simplex algorithm on 10,000 parameter sets sampled with latin hypercube sampling (with *lhs* R package [43]) and chose the one with the highest posterior value at the end of the chain. The posterior is highly multimodal and thanks to this initialization, the MCMC chain explored the region of the highest mode. The covariance matrix of the proposal distribution was initialized using adaptative MCMC as in Dureau et al. [41].

### 2.5 Model comparison

#### 2.5.1 Statistical indicators

In order to identify the best model, the Deviance Information Criterion (DIC) [44] is used. DIC is an indicator that combines a measure of model fit and a penalty on model complexity, commonly used with MCMC estimations. The best model is the one with the smallest DIC. As it does not enable comparison of models with differing number of observations, we also calculate mean RMSE (root mean square error) between observations and simulations of the model with parameters sampled in the posterior distribution (with observation indices from *t*_0_ to *T*, and *N* simulated trajectories):

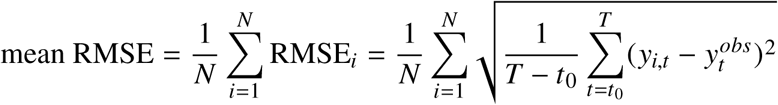

We first calculated this indicator on the data used for estimation (separating NDSS data and DENFREE data). Then we used it to assess the predictive performance of the model, comparing projections of the model with NDSS observations for 2014 and 2015.

#### 2.5.2 Epidemiological indicators

Models are also compared according to several indicators to describe their epidemiological behaviour. The basic reproduction number, the observation rate and the initial proportion of susceptibles are estimated using the MCMC chain. In the model with asymptomatic infections we report the estimated proportions of asymptomatic and hospitalized cases. With parameters sampled in the posterior distribution, we can also re-simulate the model to study hidden states, such as the susceptible and infected classes. The effective reproduction number (*R*_*e*_) is calculated as the seasonal basic reproduction number multiplied by the proportion of susceptibles at each time step, as indicated in Table 5. We then calculate the annual incidence proportion as the total number of infections over one year divided by the total population of susceptibles at the beginning of the year. In models with two strains, we separate the annual incidence of primary infection (as the total number of primary infections over one year divided by the total population of naive individuals at the beginning of the year) and secondary infection (as the total number of secondary infections over one year divided by the total population of susceptibles to one strain only at the beginning of the year).

**Table 5.** **Reproduction numbers calculation in each model.**

|  | SEIR, Laneri, SEIAR | Pandey | SEIR2 | SEIR2psi |
| --- | --- | --- | --- | --- |
| $R_0(t)$ | $\frac{\beta(t)\sigma}{(\gamma+\mu_H)(\sigma+\mu_H)}$ | $\frac{\beta_H\beta_V(t)\sigma\tau}{(\gamma+\mu_H)(\sigma+\mu_H)\mu_V(\mu_V+\tau)}$ | $\frac{\beta(t)\sigma}{(\gamma+\mu_H)(\sigma+\mu_H)}$ | $\frac{\beta(t)\sigma}{(\gamma+\mu_H)(\sigma+\mu_H)}$ |
| $R_e(t)$ (or $R_e^i, i = 1, 2$) | $R_0(t) \frac{H_S(t)}{N}$ | $R_0(t) \frac{H_S(t)}{N} v_S(t)$ | $R_0(t) \frac{H_S(t)+H_{Sj}(t)}{N}$ | $R_0(t) \frac{H_S(t)+H_{Sj}(t)}{N} \left[ \frac{H_{Ii}(t)+\psi H_{Iji}(t)}{H_{Ii}(t)+H_{Iji}(t)} \right]$ |

Calculations are made using R version 3.2.2 [45], and graphics using *ggplot2* [46].

## 3 Results

### 3.1 Statistical comparison

For single strain models, the SEIR and Pandey models proved the best with the DIC criterion (cf. Table 6). For two strain models, the SEIR2psi proved best. As regards simulation-based indicators on the 2002-2014 data, SEIR, Laneri and Pandey models have RMSE values in the same order of magnitude. Indeed, they produce a similar dynamic with respect to the 2002-2014 data (cf. Figures 7-8), with a period of approximately six years, and a large overestimation of the 2002 outbreak (the RMSE for 2002 is far higher (> 50) than the average for the other years (<20)). Due to the small number of observed asymptomatic cases, SEIAR model also produces a similar dynamic. Models with two strains outperform all the other models in terms of RMSE, but the difference is mainly explained by the first year of simulation (2002). When visualizing the simulations compared to the data (cf. Figures 7-8), all models underestimate a large number of epidemic peaks and all models but the ones with two strains overpredict the first epidemic peak (as pointed out in Table 6). SEIR2psi is the model in which the large epidemic in 2007 is best reproduced and overall, SEIR2psi is the model that reproduces most accurately the observed data.

**Table 6.** **Information criteria in deterministic model.** DIC is the Deviance Information Criterion [44]. RMSE is the root mean square error between simulations and observations: it is calculated separately on the datasets used for estimations (NDSS data for 2002-2013 and DENFREE data for 2012-2013) and on the test set (NDSS data for 2014-2015). It is also computed on separated years in order to highlight well or badly estimated years: for example, for each simulation *i*, 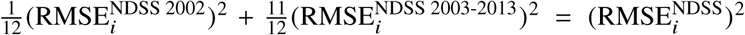. Convergence diagnosis are displayed in Appendix (trace plots et correlation plots respectively in Figures 12 and 13).

| Model | SEIR | Laneri | Pandey | SEIAR | SEIR2 | SEIR2psi |
| --- | --- | --- | --- | --- | --- | --- |
| **nb parameters** | 7 | 8 | 9 | 8 | 10 | 11 |
| **nb observations** | 656 | 656 | 656 | 686 | 686 | 686 |
| **ESTIMATION SET** |  |  |  |  |  |  |
| **DIC** | 4774 | 4818 | 4778 | 4990 | 4800 | **4604** |
| **mean RMSE NDSS** | 23.6 | 23.8 | 23.8 | 24.6 | 18.8 | **16.8** |
| *mean RMSE NDSS 2002* | *52.2* | *51.4* | *52.6* | *55.9* | *14.4* | *15.1* |
| *mean RMSE NDSS 2003-2013* | *19.0* | *19.4* | *19.2* | *19.4* | *19.1* | *17.0* |
| **mean RMSE DENFREE** | 23.7 | 23.8 | 23.0 | 24.3 | **20.3** | 20.7 |
| **TEST SET** |  |  |  |  |  |  |
| **mean RMSE 2014-2015** | 14.7 | 14.7 | 15.3 | 15.4 | 21.3 | **9.8** |
| *mean RMSE 2014* | *18.5* | *18.5* | *18.9* | *19.5* | *25.8* | *11.4* |
| *mean RMSE 2015* | *9.6* | *9.1* | *10.2* | *9.6* | *15.4* | *7.7* |

**Figure 7.**
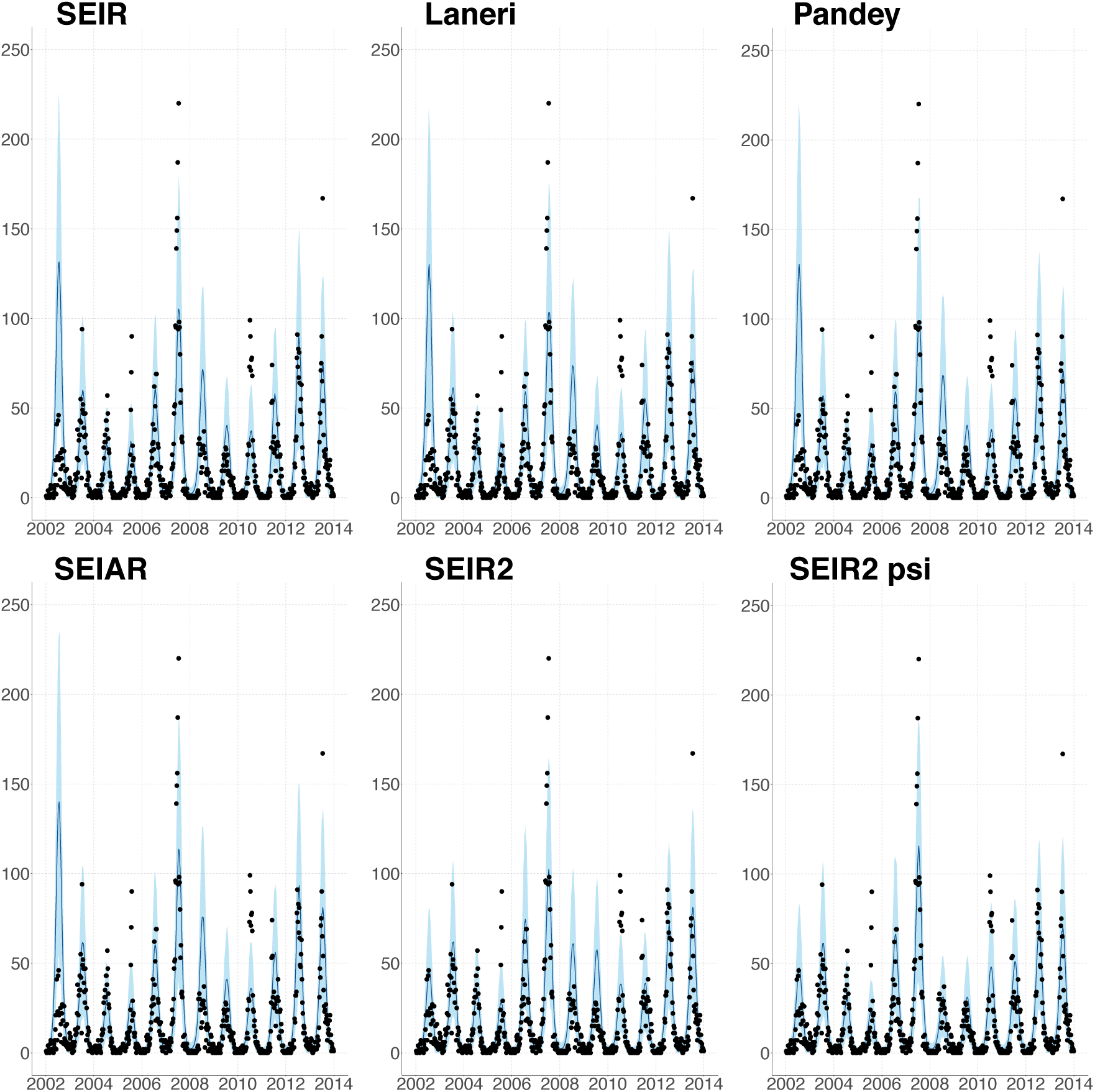
Number of observed cases per week and NDSS data, 2002-2013. Simulations with negative binomial noise using parameters from the posterior distribution (SEIR/Laneri/Pandey/SEIR2/SEIR2psi: observed NDSS cases, SEIAR: hospitalized cases). Posterior median (solid line), 95% credible intervals (shaded blue area) and NDSS data points (black dots).

**Figure 8.**
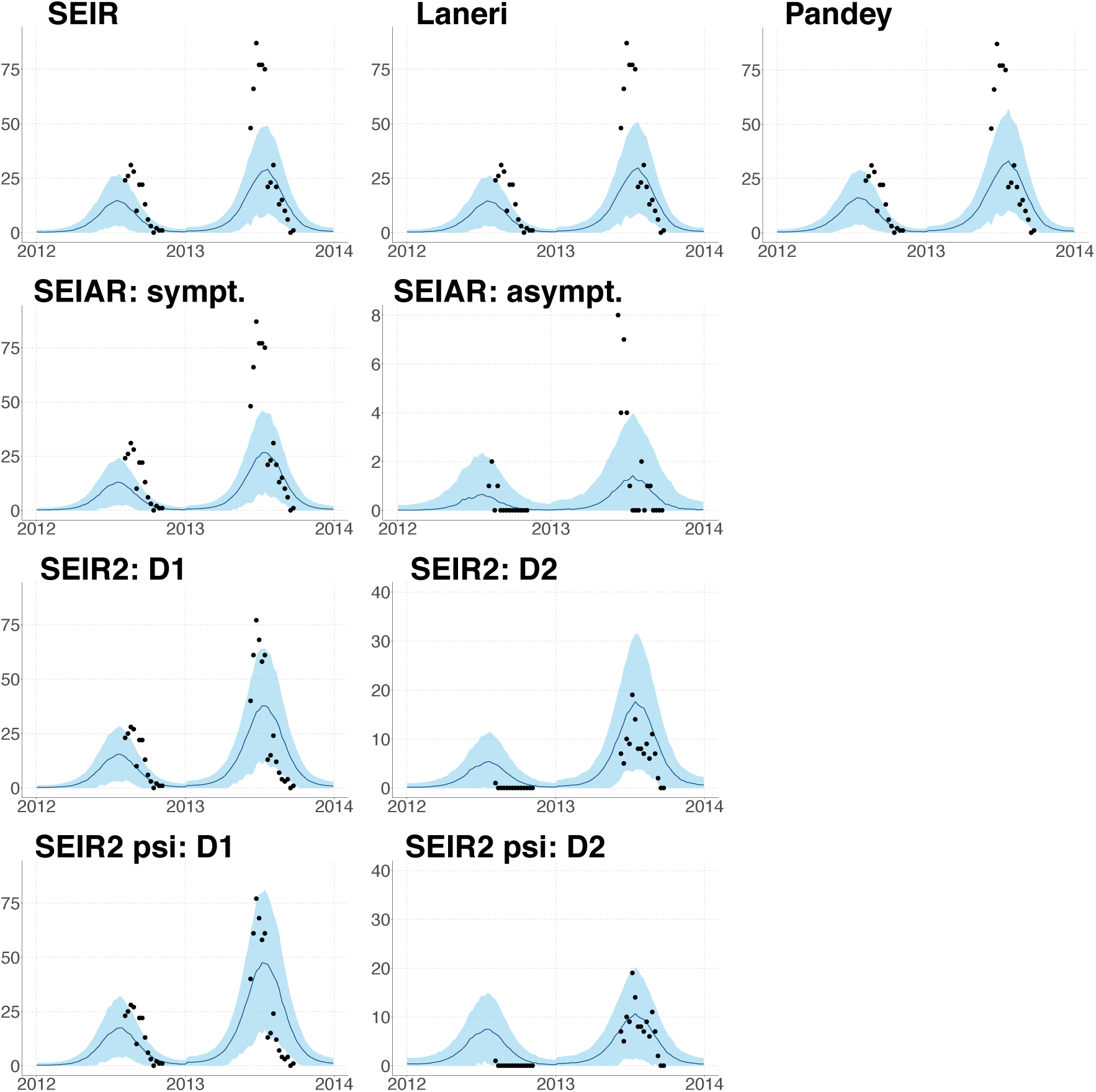
Number of observed cases per week and DENFREE data, 2012-2013. Simulations with negative binomial noise using parameters from the posterior distribution. Posterior median (solid line), 95% credible intervals (shaded blue area) and DENFREE data points (black dots).

Considering the predictive capacity of the models, only SEIR2psi model predicts a smaller epidemic in 2014, as was observed in the data (cf. Figure 9). Across the other models, the predictions for 2014-2015 are qualitatively similar: they largely overestimate the 2014 epidemic, and to a lesser extent the one in 2015.

**Figure 9.**
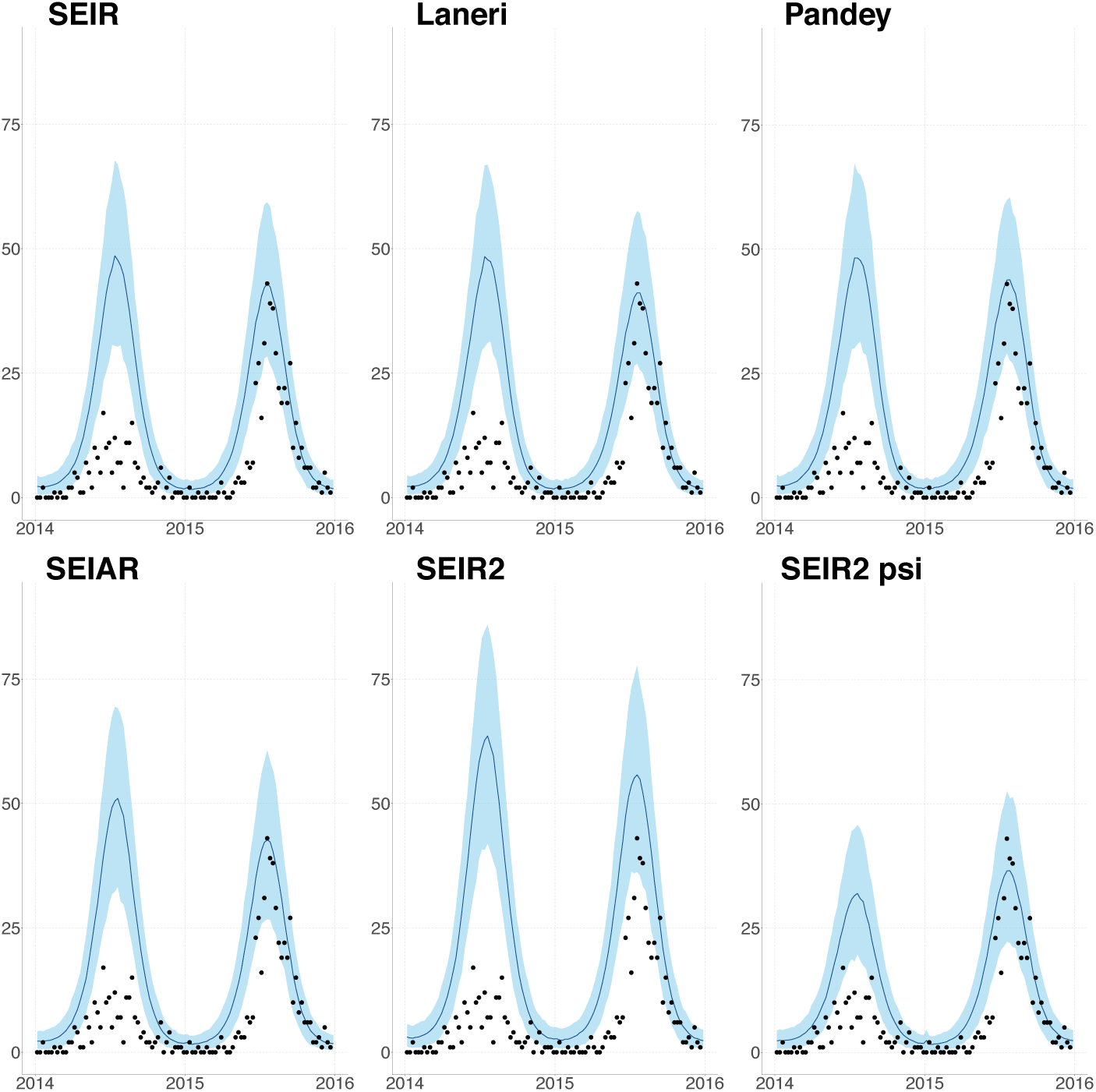
Projections of the number of observed cases per week and NDSS data, 2014-2015. Simulations with negative binomial noise using parameters from the posterior distribution (SEIR/Laneri/Pandey/SEIR2/SEIR2psi: observed NDSS cases, SEIAR: hospitalized cases). Posterior median (solid line), 95% credible intervals (shaded blue area) and NDSS data points (black dots).

### 3.2 Epidemiological comparison

The average *R*_0_ is estimated to be between 2 and 3 in most of the models and the maximum value between 3 and 4 (cf. Table 7), except with the Pandey model, in which it is higher (mean value above 3 and maximum value above 6.0). The estimated values are very close in the SEIR, Laneri and SEIAR models. The estimates are on the lower side of those estimates for South East Asia in general [47, 48, 49]. In particular, our estimates are close to the estimates for *R*_0_ in Cambodia made from age-stratified case-notification data [49], as far as the estimates based on different data types and models are comparable. Nevertheless, our estimation of *R*_0_ strongly depends on the estimation of the initial proportion of susceptibles (*H*_*S*_(0)), which is unknown in the case of Cambodia and can bias the estimates. The effective reproductive numbers (*R*_*e*_) have a mean value around 1 and a maximum value around 1.8 in most models, but it is higher in the Pandey model.

**Table 7.**
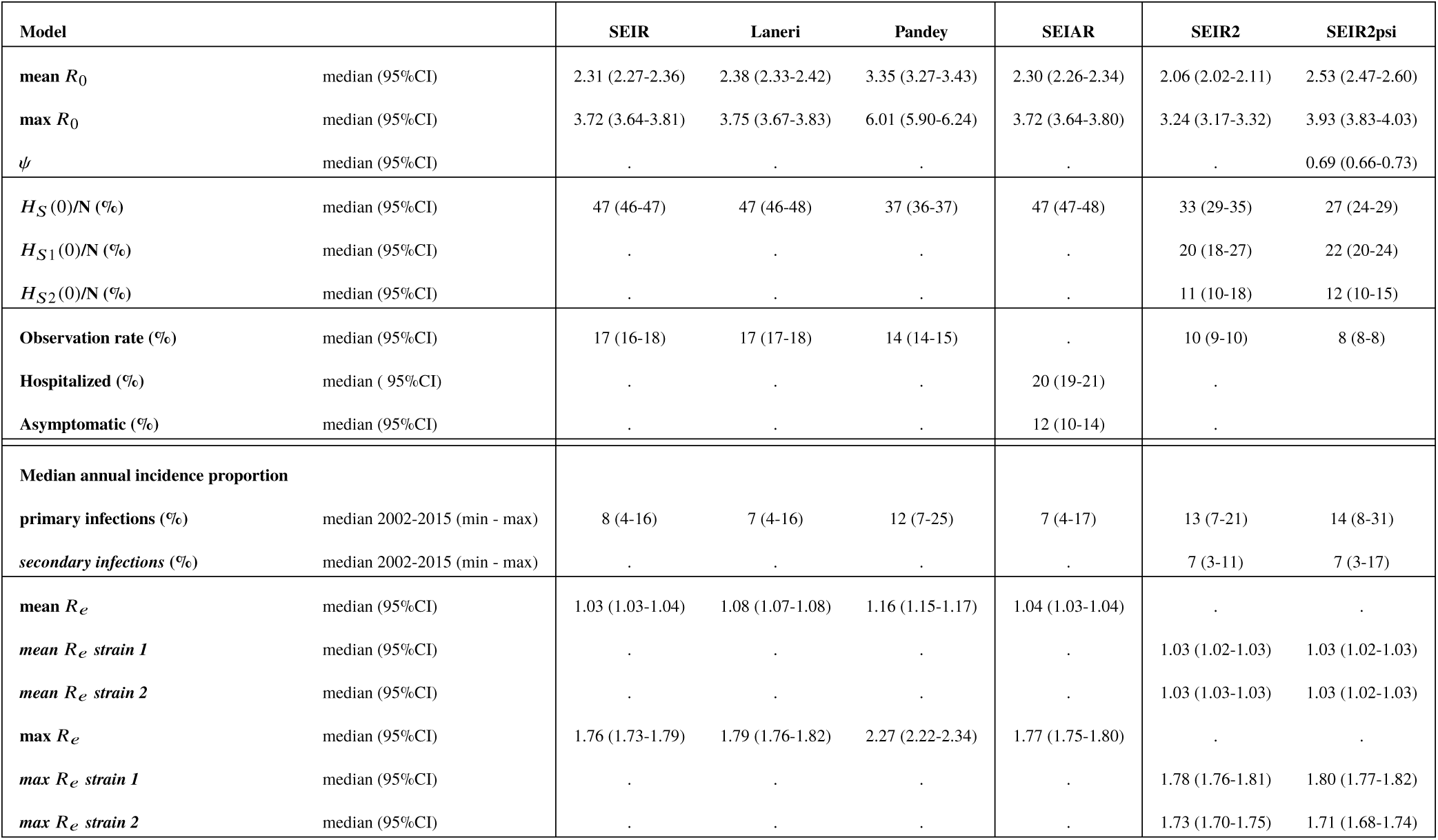
**Epidemiological criteria among children under 15 years old.** Estimated parameters from the posterior distribution and indicators based on simulations over 2002-2015.

In SEIR2psi model, the parameter *ψ* quantifying the interaction between strains is inferior to 1, suggesting a reduced infectivity of secondary infections on average, as in Aguiar et al. [36], or in Coudeville and Garnett [50]. This suggests that at the population level cross-protection is more important than cross-enhancement to explain the results observed in the field.

It is estimated in the models that approximately half of the children are susceptible to the disease, which is close to the informative prior used. This proportion is lower in the Pandey model. In the models with two strains, the number of susceptibles to both strains is smaller, but as a whole, less children are immune to the disease, since more than 30% have experienced only one infection. The proportion of children who are susceptible to one or both strains are however correlated in the MCMC chain (cf. Figure 13 in Appendix), indicating that their relative shares are not well identified. These values are in the range of the measures of seroprevalence in several Asian countries [51]. As the measures reveal large differences between countries [51, 40, 52, 53], a seroprevalence survey in Cambodia would be particularly useful to evaluate which scenario is more plausible.

The observation rate for NDSS data varies between models, from 14 to 21% in models with one strain and between 6 and 13% in the two strain models. These values indicate that a large proportion of dengue cases are not reported in national surveillance, likely reflecting mild symptoms that do not require hospitalization or misdiagnosis or misreporting.

In the SEIR2psi model, we also plotted the current number of infected individuals for each strain (cf. Figure 10). In our simulations, the first strain is responsible for large explosive outbreaks, whereas the second one has a more regular pattern over the years. Moreover, the two strains are asynchronous and each one dominates for two or three years. It is also qualitatively close to the dynamics observed in Thailand [13] or Singapore [54].

**Figure 10.**
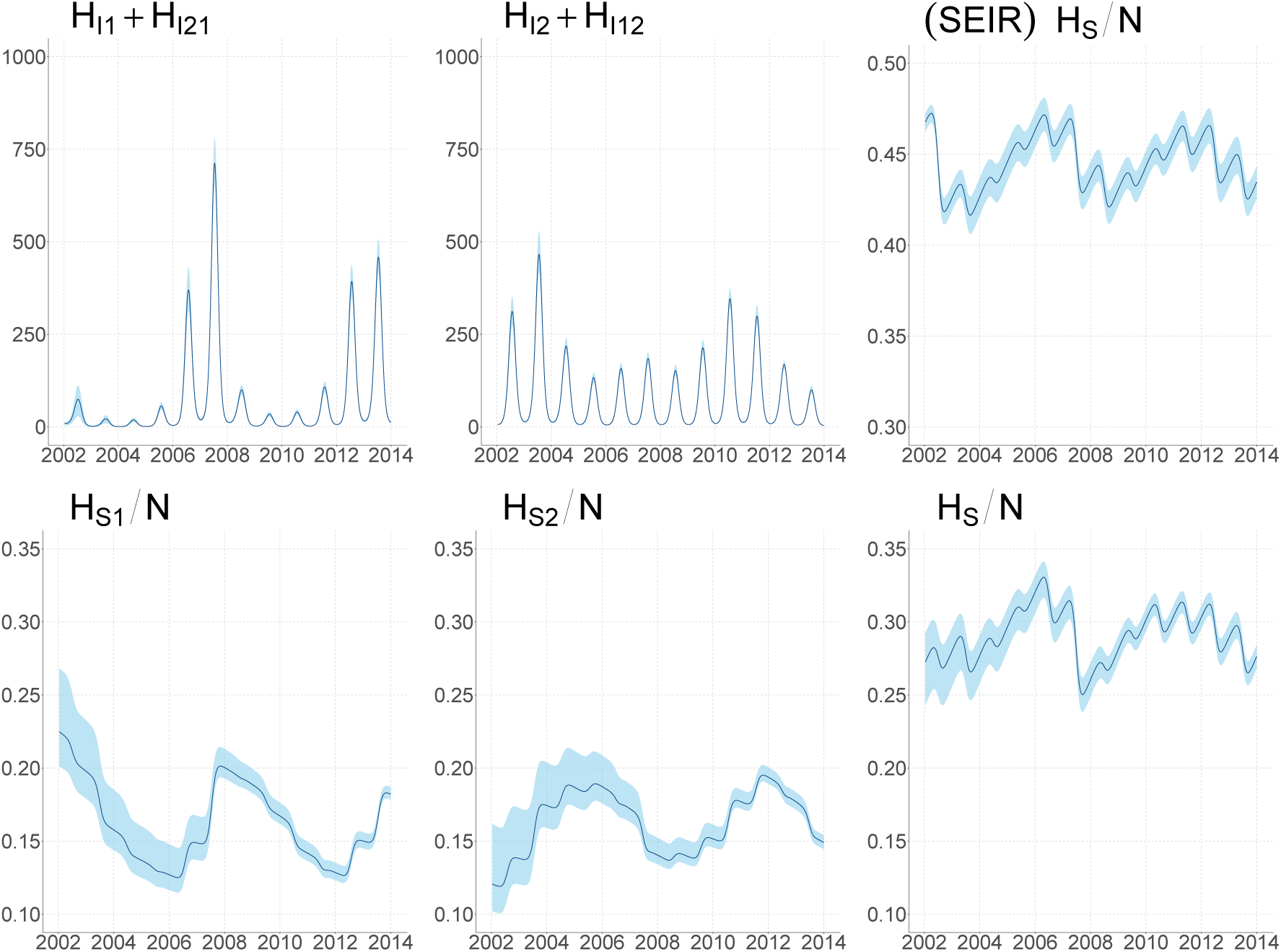
Current number of infected and proportion of susceptible individuals per serotype in SEIR2psi model. Median of the simulations and confidence intervals without observation error. For comparison, the proportion of susceptibles in SEIR model is also displayed.

The proportion of susceptible individuals also displays asynchronous dynamics between strains, as they reflect the history of past epidemics. Despite the seasonality and the year-to-year variations, the total number of susceptibles remains high (cf. Figure 10 for SEIR2psi and SEIR models), allowing the possibility for large outbreaks to occur in the future.

We calculated the annual incidence proportion as the proportion of new infections over one year among the susceptibles at the beginning of that year (cf. Figure 11). The values for primary infections are coherent with other studies in Vietnam or Indonesia who analyzed seroprevalence data or seroconversion data [40, 55, 56, 57, 52]. The incidence proportion is highly variable from one year to the next, especially in models with two strains.

**Figure 11.**
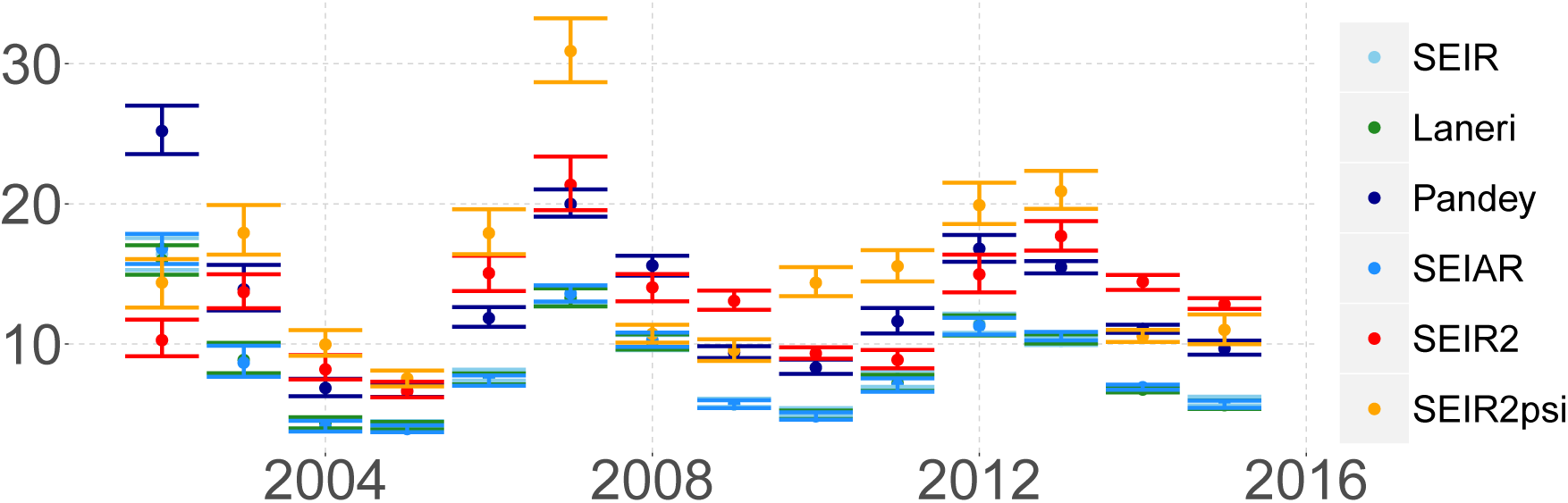
Annual incidence proportion of first dengue infection (%). Median and confidence intervals per year, based on simulations over 2002-2015.

## 4 Discussion

With two datasets reporting dengue cases in the Kampong Cham region in Cambodia, we compared several models to represent dengue transmission dynamics in a rural setting. In order to assess the quality of the models, we compared their statistical properties and their epidemiological features. In particular, in our models, the proportion of susceptibles displays year-to-year variations but remains high over the period, so that large outbreaks may occur in the future. The best model describing the dengue trend over 14 years of data was the two strain model, with reduced infectivity for secondary infected individuals. Secondary infections being more prone to severe dengue [7], these individuals may stay at home or at the hospital, and be less involved in the spread of the disease than the ones whose mild symptoms do not alter "house-to-house movement" [58, 59]. This feature was previously analytically studied and coherent with dengue incidence time series in Thailand [36, 60]. On the contrary, including vectorial transmission or a compartment for asymptomatic infections did not seem to improve the model fit despite the additional complexity. The non-utility of including vectorial components has been observed by several authors previously [29, 61, 62]. Mathematical analyses have suggested that because the time scale of the mosquito epidemiology is so fast compared to that in humans, it will be slaved by the slower human epidemiology. Thus, for understanding human disease epidemiology, mainly the dynamics of the human time scale are essential and inclusion of mosquito dynamics results in an unnecessary increase in model complexity when vector data is not available [63, 61, 62]. The main effect of mosquito dynamics is captured in our model by the seasonal forcing. The lack of additional improvement when including the asymptomatic class is likely due to the very few asymptomatic infections observed, which may be due to a very strict definition of asymptomatic infections. Yoon et al. [59] also observe many inapparent cases in their cohort but few strictly asymptomatic cases in their cluster study in Thailand. Therefore, in our model, including a compartment for asymptomatic individuals had only little influence on the overall transmission due to their small number.

We also obtained some insight on the parameters describing transmission. The average annual *R*_0_ is estimated between 2.1 and 2.6 in most models. These values are within the range observed in urban settings, suggesting that, despite very different population densities, the rural dynamics of dengue are not that dissimilar, or that dissimilarities are hidden by the variations between countries and the uncertainties due to diverse estimation procedures. The median annual incidence at primary infection over the period is between 8 and 14, with large year-to-year variations. The estimated observation rate on surveillance data varies between models, (14-21% in models with one strain and 6-13% in two strain models), indicating in both cases a high proportion of unreported infections. These values are in line with the large underrecognition highlighted in Wichmann et al. [64], and also with other studies in South-East Asia [59, 55, 57].

This work has however several limitations. Firstly, model comparison was not straightforward between one strain and two strain models, both statistically and epidemiologically. From the statistical point of view, the differing number of observations between models led us to use simulation based-indicators. From the epidemiological point of view, single strain models, two strain models and observations on four serotypes may be hard to compare because some indicators cover different interpretations. For example, in single strain models, there is no distinction in the susceptible class between individuals immune to one strain only and naive ones, and there is no strain specific *R*_0_ or incidence.

Secondly, the selected model formulations were restricted due to data availability. In particular, despite the endemicity of the four serotypes in rural Cambodia, we did not consider more than two dengue serotypes. This was done to limit model complexity, especially in the number of unknown initial conditions, but has also been previously shown to adequately describe dengue dynamics [39]. When two serotypes were considered, we tested only interactions in terms of enhancement or restriction of infectiousness. We did not include models with (temporary) cross immunity, because of the too large increase in the number of parameters with respect to the data. We also did not include models with a finer spatial scale, even if small scale transmission plays a decisive role in dengue dynamics [12, 65]. On the one hand, NDSS data were only available at the district level, which was too large to follow transmission chains and too small for observing a sufficient number of cases. On the other hand, the clustered sampling protocol in the DENFREE study abnegated the interpretation of the spatial distribution of community cases. We also restricted the analysis to children under 15 years old and did not study the role of adults in transmission.

Thirdly, the projections are not completely able to describe the observed data, as most models overestimate the dengue epidemic in 2014. Nevertheless, 2014 was a particular year, with the lowest number of cases in the whole time series, maybe due to particular climatic conditions. In many countries in South-East Asia, except Malaysia, the reported incidence was lower than in 2013 [66]. Many provinces of Thailand also reported fewer cases than usual in 2014 [67]. Our models are deterministic and do not take into account variations due to demographic stochasticity or environmental hazards such as climate.

Despite these limitations, combining two datasets permitted us to overcome some observation biases, such the fact that surveillance data did not report serotype and DENFREE data did not reflect the long term dynamics. Nevertheless, some information is lacking in both datasets, in particular that on seroprevalence. Clearly the parameter estimations depend on the immunological status at the beginning of the simulations. As in our previous work [34], our modeling study stresses the importance of seroprevalence data in order to more accurately estimate the initial conditions of our simulation and reduce identifiability problems. A seroprevalence survey in Cambodia would be of great value to evaluate the dengue burden, transmissibility potential and consider vaccination scenarios.

In conclusion, our analyses highlight the importance of using two-strain models with interacting effects and the lack of added value of incorporating vector and asymptomatic components. Although model complexity is framed by the scientific objectives, it must also be driven by the available data. The unavailability of mosquito data and the very low incidence of asymptomatic infections questions their incorporation explicitly in the models. Another important aspect is related to choosing the best model considering the available data. In quantitative epidemiology, in recent years, it has been standard to use statistical indices computed on estimated likelihood [29, 28]. In this work, we discussed not only the goodness of fit of the models, but also their prediction capacity and epidemiological features. The sole use of statistical indices is not enough and it is crucial to take into account some epidemiological features (such as *R*_0_, annual incidence, dynamics of the susceptible class) of the disease studied into account.

## Acknowledgments

All the authors have been supported by a grant from Agence Nationale de la Recherche for the PANIC project (Targeting PAthogen’s NIChe: a new approach for infectious diseases control in low-income countries: ANR-14-CE02-0015-01). The research leading to these results has also received funding from the European Commission Seventh Framework Program FP7 for the DENFREE project under Grant Agreement 282 378. CC and BC are partially supported by the "Pépiniere interdisciplinaire Eco-Evo-Devo" from the Centre National de la Recherche Scientifique (CNRS). CC is supported by the PhD studentship of the Groupe des Ecoles Nationales d’Economie et Statistique (GENES). The funders played no role in the study design, data collection, analysis, or preparation of the manuscript.

## A Data and codes

The datasets and codes supporting this article are available at https://github.com/clchampag/KC-dengue.

## B Convergence diagnosis

Figure 12 – **Trace plots for the MCMC algorithm, with 100,000 iterations.** a) SEIR. b) Laneri. c) Pandey. d) SEIAR. e) SEIR2. f) SEIR2psi

Figure 13 – **Correlation plots of the MCMC algorithm, with 100,000 iterations.** a) SEIR. b) Laneri. c) Pandey. d) SEIAR. e) SEIR2. f) SEIR2psi

